# Early developmental neuronal activity inhibits oligodendrocyte differentiation through AMPA receptor activation

**DOI:** 10.1101/2025.10.17.683139

**Authors:** Tessa Allen, Graham Peet, Luis Gomez Wulschner, Won Chan Oh, Wendy Macklin

## Abstract

Oligodendrocytes produce myelin, a lipid-rich membrane that wraps neuronal axons in the central nervous system to provide metabolic and trophic support and allow for saltatory conduction. Developmental myelination requires precisely timed and localized neuron-oligodendrocyte communication. In the mature brain, neuronal activity promotes oligodendrocyte precursor cell (OPC) proliferation and differentiation, but how OPCs respond to neuronal activity in early brain development, prior to the onset of myelination, is less well characterized. Here, we investigate how sensory-evoked and chemogenetically altered neuronal activity affect oligodendrocyte maturation in the olfactory system, somatosensory cortex, and corpus callosum in mice. We find that early neuronal activity inhibits oligodendrocyte differentiation and that reduced neuronal activity relieves this inhibition, thereby increasing OPC differentiation. Additionally, single-cell RNA sequencing revealed transcriptional changes in oligodendrocytes when neuronal activity was reduced, including upregulation of glutamate receptor gene expression. Finally, we identify AMPAR signaling as a critical regulator of oligodendrocyte differentiation ex vivo.

## Introduction

Oligodendrocytes are the myelinating cells in the central nervous system, providing neurons with metabolic and trophic support and enabling saltatory action potential propagation. Oligodendrocyte progenitor cell (OPC) differentiation and myelination during development require precise spatial and temporal regulation through bidirectional communication between OPCs and neuronal axons, yet this important developmental signaling is not well understood^1^. Several studies have shown that adult neuronal activity promotes oligodendrocyte proliferation and differentiation (oligodendrogenesis)^2^. In early development, however, when OPCs are rapidly proliferating and prior to normal developmental myelination, the relationship between oligodendrocytes and neurons remains largely unclear.

In the mouse olfactory system, the lateral olfactory tract (LOT) is one of the earliest white matter tracts to develop in the CNS, as the relay of olfactory information is critical for early mouse development^3^. Mitral cells and tufted cells in the olfactory bulb send axons to form the LOT as early as E11, and robust myelination of this tract begins around P7^4–6^. Our lab and others have shown that chronic suppression of neuronal activity in adult mouse models leads to decreased oligodendrogenesis and hypomyelination of the LOT^6,7^. Here, we use this system to address how neuronal activity influences oligodendrogenesis in early development, studying the LOT prior to active myelination.

In mouse cortical development, cortical projection neurons begin to extend axons contralaterally to form the corpus callosum around embryonic day (E) 15^8^. During the first week of postnatal development, there is extensive early axonal exuberance followed by dynamic circuit refinement^9^. In similar developmental timing, OPCs differentiate from neural stem cell-derived radial glia in the brain starting at E12.5^10^. They are abundant in the corpus callosum by postnatal day (P) 0, and begin to populate the cortex at birth^11^. In the corpus callosum, OPCs start to differentiate into premyelinating oligodendrocytes around postnatal day (P) 7, and robust myelination begins around P14^12^. Thus, during the first postnatal week, axons are highly active, yet unmyelinated. Indeed, one study showed that sensory deprivation by whisker plucking for the first five postnatal days causes increased oligodendrogenesis in the deprived region of the barrel cortex^13^. We hypothesized that before axons are physiologically ready for myelination, the influence of neuronal activity on oligodendrogenesis is different than during active myelination. The current study aims to define how neuronal activity alone regulates oligodendrogenesis before the onset of developmental myelination.

Glutamatergic signaling from neurons to OPCs and oligodendrocytes regulates oligodendroglial maturation and function in the adult central nervous system (CNS). Glutamate signals through various ionotropic and metabotropic receptors, including receptors activated by α-amino-3-hydroxy-5-methyl-4-isoxazolepropionic acid (AMPARs), which are expressed by oligodendrocytes and their precursors^14^. AMPARs are present in OPCs across species early in development, suggesting that glutamatergic signaling plays an important role in developmental oligodendrogenesis and myelination^15–17^, although their impact is debated. In vitro studies show that AMPAR stimulation inhibits OPC proliferation and differentiation^18–20^, and both in vitro and in vivo studies show it increases OPC migration^15,21,22^. Conversely, AMPAR-mediated glutamatergic signaling on OPCs stimulates oligodendrogenesis in vivo and promotes myelination in later development and adulthood^18,23,24^. An important caveat is that these published in vivo studies mostly examine glutamatergic signaling in adulthood or in late development, after robust myelination has begun. The effects of glutamatergic transmission prior to this essential developmental myelination period have not been studied; thus, how glutamatergic neuronal activity regulates OPCs and oligodendrocytes in the early developmental environment remains unclear. Here, we investigate the role of glutamatergic neuronal activity with a focus on the contribution of oligodendroglial AMPARs to OPC differentiation in early postnatal development.

## Methods

### Animals

All animal experiments were performed with approval from the Institutional Animal Care and Use Committee at University of Colorado, Anschutz Medical Campus. For DREADD experiments, *Neurod6*^*tm1(cre)Kan*^ (NexCre+/-)^25^ mice were crossed to WT *C57BL/6J* mice and genotyped. Mice were sacrificed via perfusion for immunohistochemistry (IHC) or decapitation for electrophysiology at P7 or 8 as indicated. Unilateral Naris Occlusion (UNO) mice were sacrificed via perfusion at P5. For single cell RNASeq (scRNAseq) experiments, NexCre+/-mice were crossed to WT *C57BL/6J* mice and genotyped. After DREADD/CNO exposure (see in vivo DREADD injections), P8 mice were anesthetized with isoflurane, sacrificed via decapitation, and processed for single cell analyses (see below). For ex vivo cortical slice culture experiments, NexCre+/-mice were sacrificed at P4 by decapitation. Both male and female mice were used for all in vivo and ex vivo experiments. Mouse weights were recorded daily to monitor any changes in health compared to litter matched controls, and no differences were noted (data not shown). All animals were treated and handled equivalently between experimental conditions.

### Unilateral Naris Occlusion

Unilateral naris occlusion (UNO) was performed as previously described^26^. Briefly, P0 pups were cryoanesthetized and then immediately subjected to electrocautery for 1 second on the right nostril under a dissecting microscope. During electrocautery, care was taken to avoid contact of the electrocautery unit with any non-superficial tissues. Upon perfusion, scar formation over the right nostril was confirmed.

### In vivo DREADD Injections

Intracranial injections were performed according to our previous study^27^. Briefly, adeno-associated viruses (AAVs) were injected by hand. Neonates (P0-1) were cryoanesthetized, and following cessation of movement, a solution of recombinant AAVs (7×10^1 2^ viral copies [vc]/mL) (addgene) was injected bilaterally into layer 2/3 of the somatosensory cortex using a pulled glass needle (30–40 μm)^28^. The needle was held perpendicular to the skull surface during insertion to a depth of approximately 200–300 μm. Once the needle was in place, viral solution (1 μL) was manually injected (Microdispenser, VWR) into each hemisphere. AAVs included pAAV-hSyn-DIO-hM4D(Gi)-mCherry(Gi-DREADD) (addgene) or pAAV-hSyn-DIO-hM3D(Gq)-mCherry (Gq-DREADD) (addgene). Litter and age-matched control animals (NexCre-) received the same amount of AAV solution. After injection, pups were placed on a warming pad for recovery (15 min) and returned to the home cage^29^. All experimental and control animals were handled equivalently. In vivo clozapine N-oxide (CNO) and 5-Ethynyl-2-deoxyuridine (EdU) administration Oral CNO administration was performed according to previous studies^30^. Briefly, CNO was delivered by carefully handling pups and using a pipette to feed pups orally. CNO (1 mg/kg) was administered twice daily^28,30,31^. Litter and age-matched controls received the same concentration of CNO. For EdU administration, 5 μg/g EdU was administered IP once daily from P5-P7.

### Cortical Explant Culture

300 μm thick cortical slices from P4 NexCre+/-mice were prepared^32^. Brains were dissected in carbogenated dissection media (0.5 mL 1M CaCl2, 2.5 mL 1 M MgCl2, 0.9 g Glucose, 0.149 g KCl (or 2 mL of 1M KCl), 1.09 g NaHCO3 (Sodium bicarbonate), 40 g Sucrose, 1 mL 0.5% phenol red, dH2O to 500 mL, sterile filtered) and cultured in culture media (170 mL dH2O, 2.1 g MEM, 50 mL Horse serum, 1.25 mL 200 mM L-glutamine, 0.25 mL CaCl2, 0.5 mL MgSO4, 0.58 g D-glucose, 0.11 g NaHCO3 (sodium bicarbonate), 1.79 g HEPES, 0.012 mL 25% Ascorbic acid, 0.25 mL 1 mg/mL Insulin. Osmolality between 317-323, pH 7.28, Sterile filtered). Culture media was changed every 48 hours until 6DIV when treatment was started. For tetrodotoxin (TTX) experiments, 1 μM TTX in media or media alone was bath applied to the slices for 48hrs. The slices were then fixed in 4% paraformaldehyde (PFA) in phosphate buffered saline (PBS) for 20 minutes and stained for oligodendrocyte lineage markers. For DREADD experiments, immediately after plating from P4 brains, 1μL Gi-DREADD or control pAAV-hSyn-mCherry (mCh) was pipetted on the surface of each slice. At 6DIV, 1μM CNO alone in culture media; 1μM CNO + 100μM NBQX in culture media or 1μM CNO + 50μM AMPA in culture media was bath applied to the appropriate slices. Doses of culture media with 100μL 1μM CNO alone; 1μM CNO + 100μM NBQX or 1μM CNO + 50μM AMPA were spiked in at 24hrs. At 48hrs, slices were fixed in 4% PFA/PBS for 20 minutes and stained for oligodendrocyte lineage markers.

### Acute slice electrophysiology

Mice were sacrificed by decapitation at P7-8. The brain was removed from the skull and rapidly placed in ice-cold cutting solution containing (in mM): 215 sucrose, 20 glucose, 26 NaHCO_3_, 4 MgCl_2_, 4 MgSO_4_, 1.6 NaH_2_PO_4_, 1 CaCl_2,_ and 2.5 KCl. Cortical slices (300 μm thick) were prepared using a VT1000S (Leica) vibrating microtome. Slices were incubated at 32 °C for 30 minutes in a holding chamber containing 50% cutting solution and 50% artificial cerebrospinal fluid (ACSF) containing (in mM): 127 NaCl, 25 NaHCO_3_, 25 D-glucose, 2.5 KCl, 1.25 NaH_2_PO_4_, 2 CaCl_2_, and 1 MgCl_2_. After 30 min, this solution was replaced with ACSF at room temperature. Slices were allowed to recover for at least 1 h in ACSF before recording. After recovery, slices were transferred to a submersion type, temperature-controlled recording chamber (TC-324C, Warner Instruments) and perfused with ACSF. All solutions were equilibrated for at least 30 min with 95% O_2_ / 5% CO_2_^27^. Whole-cell recordings (electrode resistance, 5–8 MΩ; series resistance, 20–40 MΩ) were performed at 30 °C on visually identified, mCherry+ (Gi or Gq) cortical layer 2/3 pyramidal neurons within 40 μm of the slice surface of acute slices (P7-8) using a MultiClamp 700B amplifier (Molecular Devices). To determine the effect of short-term bath application of Clozapine N-Oxide (CNO, < 60 mins) on neuronal excitability following DREADDs expression, resting membrane potentials were recorded in current-clamp mode using potassium-based internal solution (in mM: 136 K-gluconate, 10 HEPES, 17.5 KCl, 9 NaCl, 1 MgCl_2_, 4 Na_2_-ATP, 0.4 Na-GTP, and ∼300 mOsm, ∼pH 7.26) at 30 °C in ACSF containing 2 mM Ca^2+^ and 1 mM Mg^2+27^. Additionally, rheobase of mCherry+ neurons was analyzed in response to 300-msec step current injections (5-pA increments) before and after bath application of 1 μM CNO.

### Immunohistochemistry

Mice were given an intraperitoneal injection of Fatal Plus (Vortech Pharmaceuticals). Intracardial perfusion was performed using PBS followed by 4% PFA/PBS. Tissues were dissected and then post-fixed in 4% PFA/PBS overnight and transferred to cryoprotect solution (30% sucrose in PBS). Brains were cryosectioned into 30 μm free-floating sections and stored at 4°C in 0.1% Sodium azide solution (Azide and PBS). Before staining, antigen retrieval was performed in 10 mM sodium citrate (pH 6.0) for 10 minutes at 65°C (550W), in a Pelco Biowave Pro. Sections were permeabilized in 0.3% Triton-X in PBS for 1hr at room temperature. All primary antibodies were incubated for three days at 4°C in 0.1% Triton-X in PBS. The following antibodies and concentrations were used: PDGFRα (R&D Systems AF1062) 1:500; BCAS1 (Santa Cruz sc-136342) 1:500; CC1 (Calbiochem OP80) 1:500; Olig2 (Millipore MABN50) 1:500; mCh (Aves mCherry-0100) 1:1000; PLP1 (AA3 rat monoclonal)^33^ 1:1000. Sections were rinsed for 15 minutes in PBS and then transferred to secondary antibodies. All secondary antibodies were used at a concentration of 1:1000 in 0.1% Triton-X in PBS and incubated 2 hrs at room temperature. Sections were rinsed in PBS and then mounted and cover-slipped with Fluoromount-G (ThermoFisher). For the lateral olfactory tract (LOT), images of the entire tract were tiled. For the olfactory bulb (OB), images of the entire OB were tiled. For the cortex, sample images were captured in cortical layer 4 (L4), no further than 500 μm from midline. For the corpus callosum, images were captured at midline. At least three sections per mouse were imaged and analyzed.

### Image acquisition and analysis

Fluorescent images were taken on a Leica DMi8 Stellaris 5 confocal microscope using a 40X oil immersion objective. For each antibody panel, all microscopy settings were kept constant to maintain consistent and comparable images. Cell counts using fluorescent images were collected by blinded investigators and quantified manually. Intensities of each channel were kept constant across animals to maintain consistent identification of positive cells. 3 sections from each animal were averaged.

### Single-cell RNA sequencing

Oligodendrocyte lineage cells were collected from P8 Gi-DREADD injected and CNO treated NexCre+/ -and WT controls^34^. Briefly, the cortex and corpus callosum were dissected and pooled from three NexCre+/ -and two WT littermate P8 pups that were equally treated with Gi-DREADD and CNO as above. The brains were minced and cells dissociated using papain (Worthington Biochemical). Oligodendrocytes were isolated using PDGFRα and O4-positive selection magnetic microbeads (Miltenyi). Transcriptomic libraries were captured using the 10X Chromium controller using V3 chemistry with 3’ selection (10X Genomics). Samples were sequenced to a depth of 50,000 reads per cell by the CU Anschutz Genomics Shared Resource. Sample demultiplexing, barcode processing, and alignment were performed using Cell Ranger (10x Genomics). Processed data were analyzed in R using Seurat version 5.1.0^35,36^. For initial quality control, cells containing fewer than 3000 detected genes and more than 10% percent mitochondrial transcripts were excluded from subsequent analysis. The contribution of the percentage mitochondrial gene reads to variance between cells was regressed out during data normalization and scaling. Unsupervised clustering was performed using a resolution of 0.2 in a UMAP embedding space, also used for visualization. To annotate clusters, the differentially expressed gene lists resulting from the Wilcoxon rank sum test were manually inspected, and names were assigned based on gene enrichment per cluster. Differential expression analysis across clusters was performed using a multiple comparisons-adjusted Wilcoxon rank sum test. Pseudotime reconstruction was performed using Monocle3^37^. Pathway analysis was performed using the SCPA package using Gene Ontology Biological Process term lists^38^.

## Results

### Early developmental decrease in neuronal activity in the olfactory system increases oligodendrocyte differentiation

Unilateral naris occlusion (UNO), where one nostril is permanently occluded, is a well-established model of reduced neuronal activity in the olfactory system^39^. UNO was induced at P0 to study the effect of reduced neuronal activity on oligodendrocyte development in the LOT at P5 (Fig. 1A). To confirm successful occlusion, cFos expression was assessed in the mitral and tufted cell layer of the olfactory bulb (Fig. 1B). Since mitral and tufted cells do not express NeuN but are stained by the pan-neuronal marker NISSL, we quantified cFos+ NISSL+ NeuN-active mitral/tufted cells (Figure 1C-D). NeuN-neurons in the mitral and tufted cell layer showed reduced cFos expression on the occluded side, confirming decreased neuronal activity following UNO.

**Figure 1:**
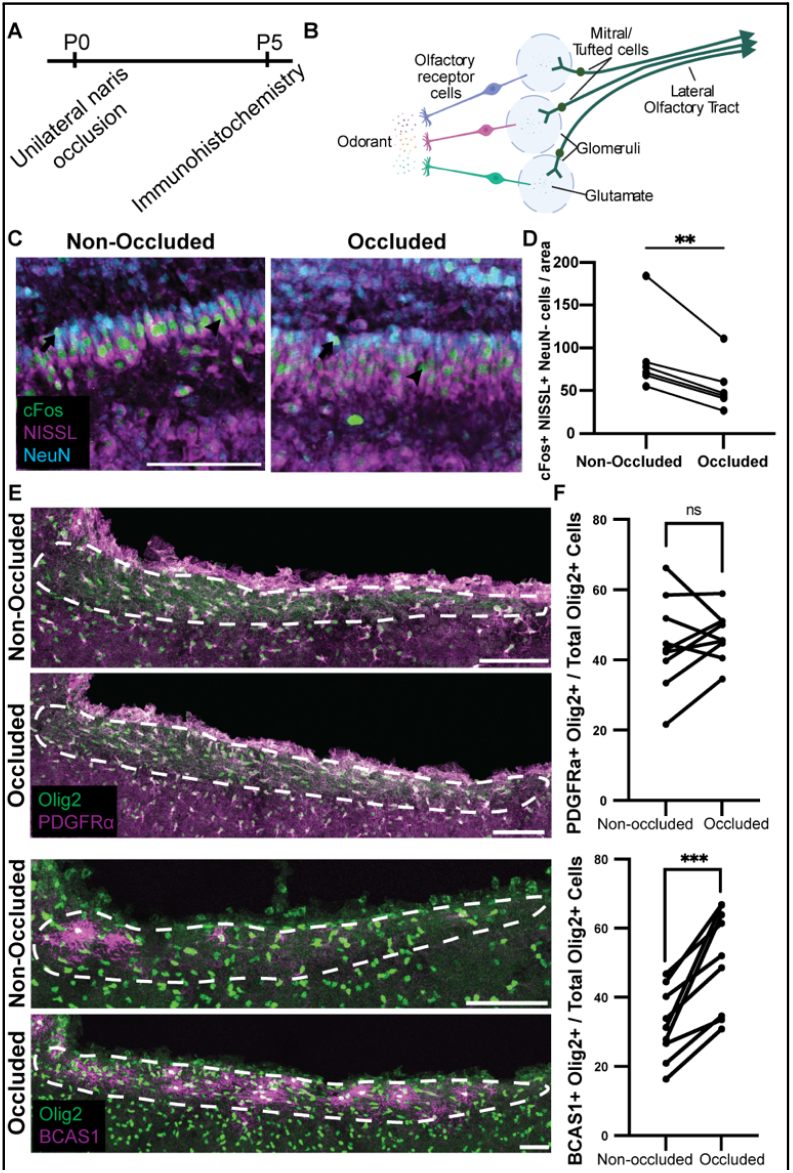
Unilateral naris occlusion causes increased differentiation of oligodendrocytes in the lateral olfactory tract. (A) Experimental timeline. P0 animals underwent UNO and were analyzed by immunohistochemistry at P5 (B) Schematic of olfactory system (C) Validation of occlusion with cFos at P5. Images are from the tufted/mitral cell layer where cells that were negative for NeuN, positive for NISSL, and positive for cFos (arrowhead) were analyzed. Cells positive for NeuN (arrow) were not analyzed as mitral and tufted cells do not express NeuN. Scale bar is 100 μm (D) Analysis of cFos+ mitral and tufted cells from occluded and non-occluded olfactory bulbs (E) Immunostaining of oligodendrocyte lineage markers Olig2, PDGFR α, and BCAS1. Dotted outline represents LOT (F) Quantification of PDGFRo and BCAS1 as a percentage of total Olig2+cells in the LOT. Scale bar is 100 μm. Data from three sections each were averaged per animal. n= 9 mice Paired Student’s t-test. ns = not significant, **p<0.01,*** p<0.001

Oligodendrocyte lineage cells in the occluded (ipsilateral) and non-occluded (contralateral) LOT were quantified using PDGFRα to visualize OPCs and BCAS1 for premyelinating oligodendrocytes. Although decreased neuronal activity did not affect the number of OPCs, surprisingly, the occluded LOT exhibited an increase in BCAS1+ premyelinating cells compared to the non-occluded LOT (Figure 1E-F). These results indicate that decreased neuronal activity increased oligodendrocyte differentiation, supporting the idea that normal neuronal activity at this developmental stage acts to inhibit OPC differentiation.

### Chemogenetic reduction of glutamatergic neuronal activity increases oligodendrocyte differentiation, but not proliferation

To investigate if this apparent inhibitory impact of neuronal activity on early OPC differentiation occurs elsewhere in the nervous system, we studied the developing somatosensory cortex and its axonal projections through the corpus callosum. In the somatosensory cortex, glutamatergic pyramidal neurons from layer II/III and V project to the same layers in the opposite hemisphere via the corpus callosum^40,41^. Importantly, during early postnatal development, there is extensive early axonal exuberance followed by dynamic circuit refinement^9^. In the corpus callosum, OPCs start to differentiate into premyelinating oligodendrocytes around P7 and robust myelination begins around P14^12^. Thus, the first week of postnatal development provides a unique environment for studying how axonal development and activity influences early oligodendrogenesis prior to the onset of myelination in the cortex and corpus callosum.

We used chemogenetics to reduce glutamatergic neuronal activity in the somatosensory cortex from P2-P8. The AAV vector encoding inhibitory designer receptors exclusively activated by designer drugs (DREADDs), AAV-hSyn-DIO-hM4D(Gi)-mCherry (Gi-DREADD), was transduced into *Neurod6*^*tm1(cre)Kan*^ (NexCre)^25^ cortex at P0 and mice were treated with the designer drug, clozapine N-oxide (CNO) from P2-P8 to reduce depolarizing activity in somatosensory cortical neurons (Figure 2A). After injecting AAV-DREADD at P0, high mCherry expression was noted in neuronal soma in cortical L2/3 and L4 of the somatosensory cortex, along with high mCherry expression in axons in the corpus callosum at P8 (Sup. Fig. 1). Decrease in resting membrane potential (RMP) and increase in rheobase in the presence of CNO (1μM) was confirmed in mCh-expressing cortical neurons from NexCre+ mice by electrophysiology (Figure 2B). OPCs (PDGFRα-positive), premyelinating cells (BCAS1-positive), and maturing oligodendrocytes (CC1-positive) were quantified and, consistent with the UNO results, no difference in OPC cell numbers was found in either the corpus callosum or the cortex. Notably, we found an increase in premyelinating cells in both the corpus callosum and cortex and an increase in maturing cells in the cortex, showing increased differentiation of oligodendrocytes when neuronal activity was chronically reduced (Figure 2C-D). This suggests that, prior to the onset of myelination, neuronal activity acts as an inhibitor of oligodendrocyte differentiation. Surprisingly, no change in OPC proliferation was seen (Sup. Fig. 2), suggesting that at these early time points, neuronal activity does not influence OPC proliferation. Together, these data show decreased neuronal activity drives increased oligodendrocyte maturation.

**Figure 2:**
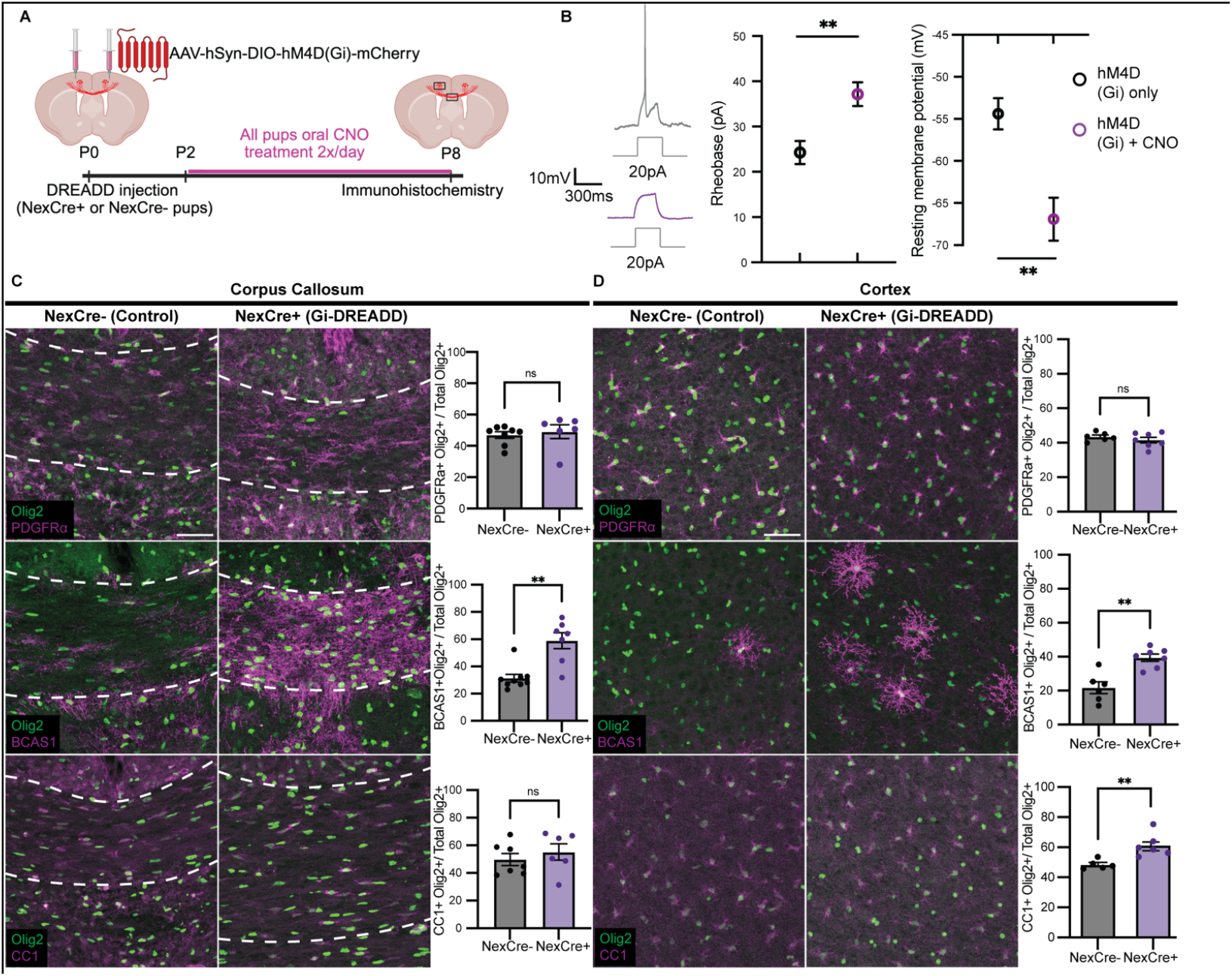
Decreased neuronal activity drives OPC differentiation. (A) NexCre+ and NexCre- control litter mates were injected at P0 or P1 with AAV-hSyn-DIO-hM4D(Gi)-mCherry (Gi-DREADD), and orally administered CNO twice daily from P2-P8. (B) Acute slices were taken from NexCre+ P6 or P7 pups and whole-cell patch clamp recordings were performed from mCherry-expressing neurons. RMP and rheobase were recorded with and without CNO in the media, n = 7 neurons from two mice (C) Representative images and quantification of the corpus callosum and (D) Somatosensory cortex show Olig2 along with PDGFRα (OPCs), BCAS1 (premyelinating) and CC1 (maturing). Data from three sections each were averaged per animal. n= 7 NexCre- control mice and 8 NexCre+ mice. Scale bar is 50um. Student’s t-test. ns = not significant* p<0.05, ** p<0.01, **** p<0.0001

### Increased excitation of glutamatergic neurons decreases oligodendrocyte differentiation, but not proliferation

To directly confirm our hypothesis that neuronal activity negatively regulates oligodendrocyte differentiation in early postnatal development, we transduced the excitatory DREADD, AAV-hSyn-DIO-hM3D(Gq)-mCherry (Gq-DREADD), into Nex-Cre cortical neurons at P0 and activated it with CNO from P2-P8 (Figure 3A). Slice electrophysiology confirmed that Gq-DREADD-expressing neurons exhibited an increased RMP and decreased rheobase upon bath application of CNO (Figure 3B).

**Figure 3:**
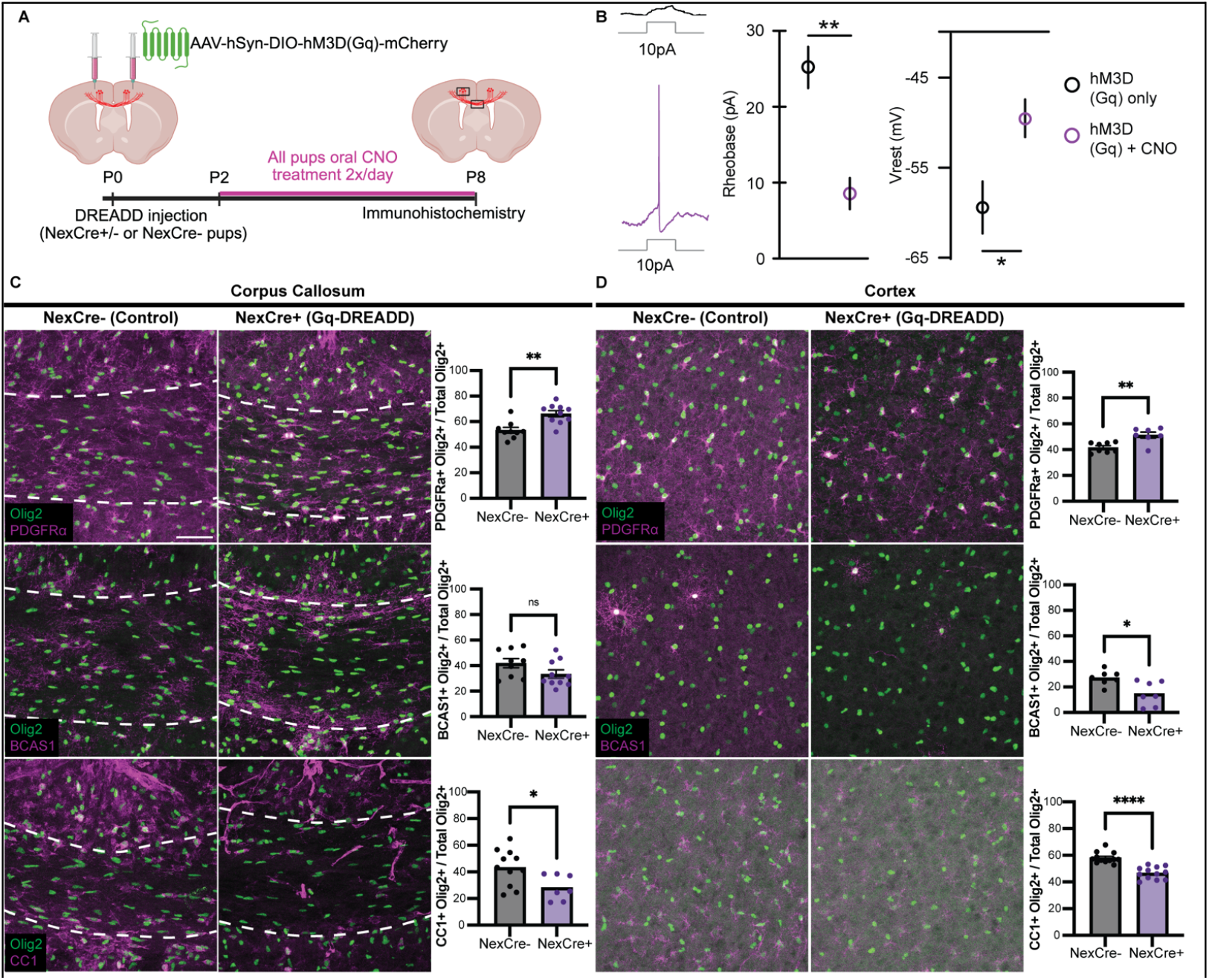
Increased neuronal activity decreases oligodendrocyte differentiation. (A) Experimental design: NexCre+ and NexCre- control littermates were injected at P0 or P1 with pAAV-hSyn-DIO-hM3D(Gq)-mCherry. and orally administered CNO twice daily from P2-P8. (B) Acute slices were taken from NexCre+ pups and whole-cell patch clamp recordings were performed from mCherry-expressing neurons. Representative traces show voltage response to 10pA injection with and without CNO. Rheobase and resting membrane potential (Vrest) were recorded with and without CNO in the media. Data is from 7 neurons from 2 pups P7 and P8 (C) Representative images and quantification of the corpus callosum and (D) Somatosensory cortex show Olig2 along with PDGFR α (OPCs), BCAS1 (premyelinating) and CC1 (maturing). Scale bar is 50um. Data from three sections each were averaged per animal. n= 7-11 NexCre- control mice and 7-12 NexCre+ mice. Student’s t-test. ns = not significant * p<0.05, ** p<0.01, **** p<0.0001

OPCs, premyelinating oligodendrocytes, and maturing oligodendrocytes were again quantified. Excitingly, increased neuronal activity reduced the number of BCAS1+ premyelinating cells in the cortex and CC1+ maturing cells in the corpus callosum and cortex (Figure 3C-D). These data further confirm our hypothesis that neuronal activity is a negative regulator of oligodendrocyte maturation in early development. Surprisingly, PDGFRα+ OPCs in both the corpus callosum and cortex were increased in response to increased neuronal activity. Similar to the Gi-DREADD experiment, there was no change in OPC proliferation (Sup. Fig2); thus, the increase in OPCs could be due to an accumulation of OPCs that are inhibited from differentiating. This further suggests that neuronal activity does not regulate proliferation at these early developmental time points, rather it independently affects differentiation, supporting our hypothesis that excitatory neuronal activity inhibits oligodendrocyte maturation prior to the onset of myelination.

### Oligodendroglial glutamate receptor expression is altered in response to reduced neuronal activity

To investigate the impact of decreased neuronal activity on early oligodendrocyte gene expression, oligodendrocyte lineage cells from Gi-DREADD-transduced, CNO-treated mice (as in Figure 2A) were isolated using magnetic bead separation and analyzed by single-cell RNA sequencing (see methods). Unsupervised clustering analysis showed successful isolation of OPCs and oligodendrocytes along a continuum of differentiation states (Figure 4A). To understand changes along this continuum, a pseudotime analysis was performed, which defined a path of state transitions between OPCs and oligodendrocytes (Figure 4B). The distribution of cells in the NexCre+ group was shifted towards more differentiated states compared to the control group (Figure 4C), which is consistent with our in vivo data showing increased oligodendrocyte maturation in response to reduced neuronal activity (Figure 2).

**Figure 4:**
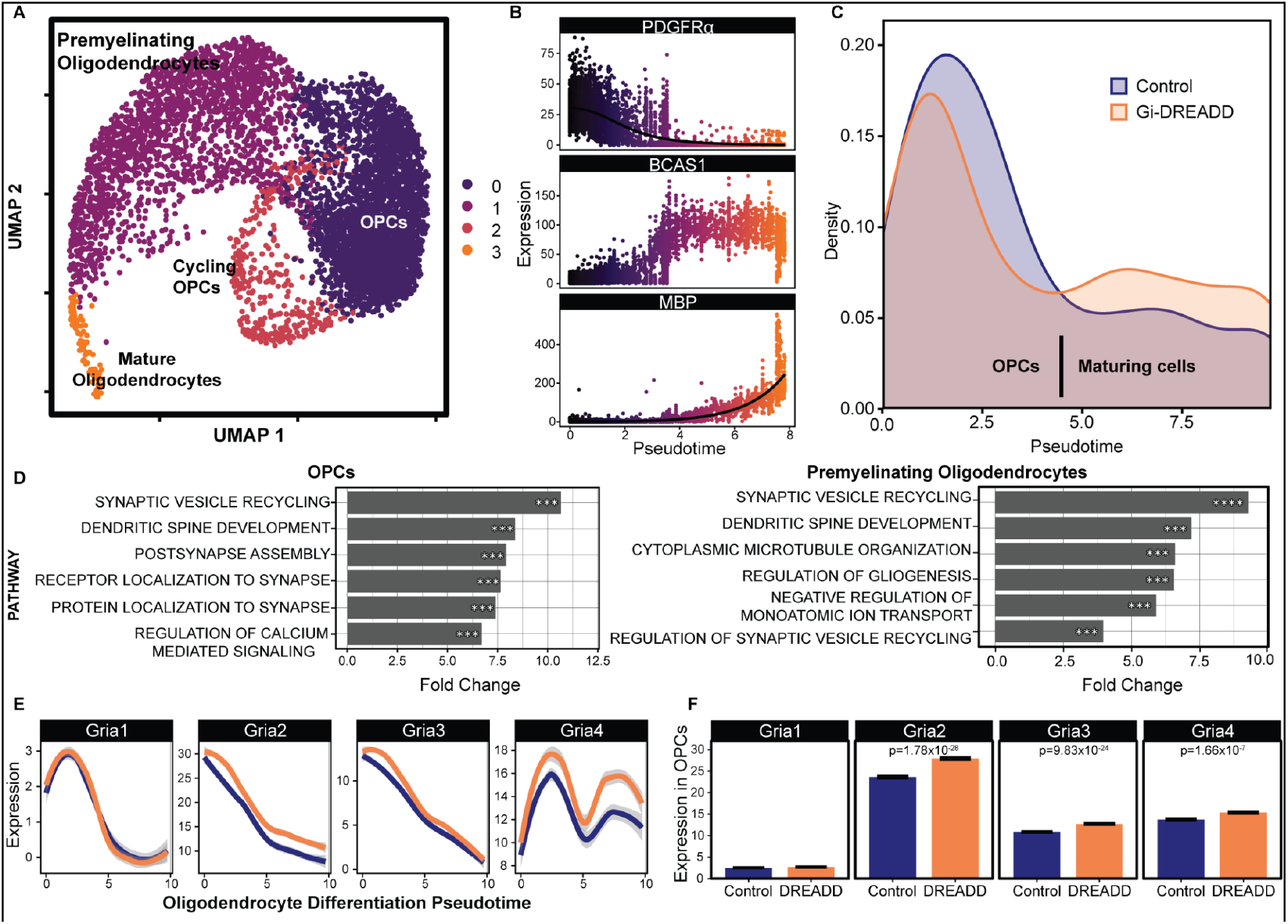
Decreased neuronal activity changes oligodendrocyte lineage gene expression and causes upregulation of AMPAR genes. NexCre- (Control, blue) and NexCre+ (DREADD, orange) were injected at P0 or P1 with AAV-hSyn-DIO-hM4D(Gi)-mCherry, and orally administered CNO twice daily from P2-P8. Brains from 3 animals each were pooled and homogenized. Oligodendrocytes were separated for sequencing by magnetic beads conjugated to PDGFRo and 04. scRNAseq analysis reveal transcriptional changes in oligodendrocyte lineage cells. (A) UMAP embedding showing oligodendrocyte lineage cells from DREADD (n=3 mice pooled) and control (n=2 mice pooled) (peach cluster: Cycling OPCs, purple cluster: OPCs, maroon cluster: premyelinating oligodendrocytes, orange cluster: mature oligodendrocytes). Refer to supplemental figure 5 for more detail (QC). (B) Oligodendrocyte lineage markers across a pseudotime cell state trajectory (Pdgfr α+ OPCs pseudotime<4, Bcas1+ premyelinating cells pseudotime >4 and <6, Mbp+ mature cells pseudotime >6). (C) Kernel density estimate of DREADD treated and control cells across pseudotime showing altered distribution of cell states. (D) Differential pathway analysis for OPC cluster (left) and premyelinating cell clusters (right) revealed synapse development and maitnence pathways. Bonferroni adjusted P values *** p < 1×10^−20^, p < 6×1 O^−20^. Wilcoxon rank sum test. (E) Expression of GluR genes across differentiation pseudotime (Control: blue, DREADD: orange) (F) AMPAR gene expression in OPCs. Bonferroni adjusted P values. Wilcoxon rank sum test. Extended data table 1 for details.

Pathway analysis revealed a variety of upregulated pathways in OPCs and premyelinating oligodendrocytes, including pathways related to postsynapse remodeling (Figure 4D). These data suggest that reduced glutamatergic neuronal activity regulates neuron-oligodendrocyte synapses in both OPCs and premyelinating cells. A variety of differentially expressed genes in response to decreased neuronal activity were identified, but interestingly, the expression of several glutamate receptor genes was significantly increased in the Gi-DREADD condition (Fig. 4E-F), again suggesting the importance of synaptic communication between neurons and oligodendrocytes. Notably, the AMPA receptor gene, *Gria2* was upregulated across the entire oligodendrocyte lineage, and *Gria3*, and *Gria4* were upregulated in OPCs (Figure 4E-F). These data show significant expression of AMPA receptor genes in oligodendrocytes at these early developmental time points, and that their expression changes in response to neuronal activity, with upregulation when neuronal activity is reduced. This suggests an important role for oligodendroglial AMPA receptors in regulating oligodendrocyte maturation in early development.

### AMPAR stimulation rescues inhibitory effects of neuronal activity on oligodendrocyte differentiation

To directly test whether oligodendroglial AMPA receptors mediate the activity-dependent regulation of oligodendrocyte maturation, we next turned to an ex vivo cortical slice culture system that allowed pharmacological manipulation of AMPAR. We first established a timeline for oligodendrocyte development ex vivo in slices from P4 mice and found peak differentiation between 6 and 8 days in vitro (DIV) (Figure 5A). To confirm that neuronal activity inhibits oligodendrocyte differentiation ex vivo, tetrodotoxin (TTX), a Na ^+^ channel inhibitor, was applied for 48 hrs. This treatment increased the proportion of cells positive for the mature oligodendrocyte marker, PLP (Sup. Fig. 3). Furthermore, pharmacological inhibition of AMPARs with NBQX for 48 hrs similarly promoted oligodendrocyte differentiation, indicating a critical role for AMPAR signaling in this process (Figure 5C). Interestingly, there was no change in the proportion of OPCs or premyelinating cells, only the increase in mature cells, suggesting that OPCs are able to differentiate to maturity within 48 hrs.

**Figure 5:**
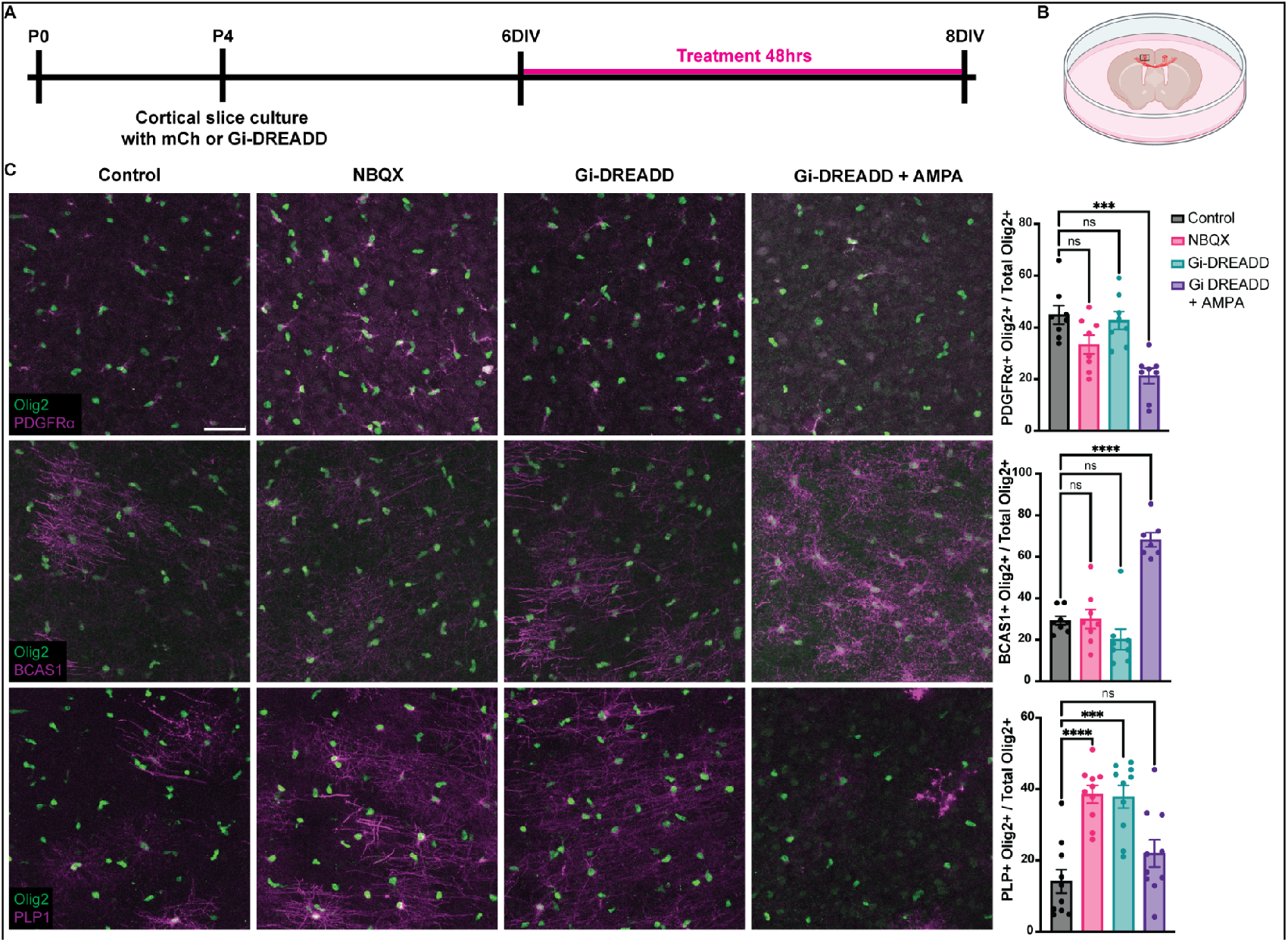
AMPA receptors regulate oligodendrocyte differentiation in organotypic cortical slice culture. (A) Schematic of experimental design. NexCre+ pups were taken at P4, and 300um thick slices of cortex were plated on a membrane with Gi-DREADD or pAAV-hSyn-DIO-mCherry (mCh). At 6 and 7DIV, 1uM CNO alone (mCh and Gi-DREADD) or 1uM CNO+ 100uM NBQX (NBQX) or 1uM CNO + 50uM AMPA (Gi-DREADD + AMPA) was bath applied to slices, and slices were fixed and stained 48hrs at8DIV. (B) Schematic of slice culture in dish Black square represents area where images were taken. (C) Images and quantification of PDGFR α (OPCs). BCAS1 (pre-myelinating oligodendrocytes) and PLP (maturing oligodendrocytes) in each experimental condition. Scale bar is 100um. n=10 mice. Student’s t-test. ns = not significant, *** p<0.001, **** p<0.0001

Finally, to determine whether AMPAR stimulation can rescue the effect of decreased neuronal activity, Gi-DREADD was used to chemogenetically reduce neuronal activity in glutamatergic projection neurons in cortical slice culture, after confirming that CNO treatment alone did not affect oligodendrocyte differentiation (sup. Fig. 4). As in vivo, Gi-DREADD activation with CNO for 48hrs increased oligodendrocyte differentiation, shown by an increase in PLP+ mature cells (Figure 5C). Surprisingly, no change in the proportion of OPCs or premyelinating cells was observed, again suggesting that OPCs can differentiate to maturity in 48 hrs (Figure 5C). To address the contribution of AMPARs to this phenotype, Gi-DREADD-expressing explants were exposed to CNO plus AMPA to reduce neuronal activity while simultaneously stimulating AMPARs. Reduced neuronal activity combined with AMPAR stimulation partially rescued the differentiation effect, as the proportion of mature, PLP+ oligodendrocytes returned to control levels (Figure 5C). Unexpectedly, this also caused a decrease in PDGFRα+ OPCs and a dramatic increase in BCAS1+ premyelinating cells (Figure 5C). These data suggest that neuronal activity acts through AMPARs to regulate the late steps of oligodendrocyte differentiation to mature oligodendrocytes, but not the initiation of OPC differentiation to premyelinating cells.

## Discussion

Here we address how neuronal activity influences oligodendrogenesis in early mouse development and find a novel role for neuronal activity in OPC differentiation: prior to the onset of developmental myelination, neuronal activity is inhibitory to oligodendrocyte differentiation and this is mediated in part by oligodendroglial AMPA receptors. Few studies have investigated this developmental question, also finding that reduced neuronal activity from P0-P6 increased OPC proliferation and oligodendrocyte differentiation in the barrel cortex^13^. With a similar sensory deprivation approach using UNO, we show that decreased activity of mitral and tufted cells in the olfactory bulb projecting through the LOT (subsequently a highly myelinated white matter tract) causes an increase in OPC differentiation in the LOT (Figure 1). This contrasts with studies in the adult olfactory system including from our group, using UNO to address the regulation of myelination^7,42^, which found decreased myelination in response to reduced neuronal activity. Our current findings are consistent with the Mangin et al (2012) study, pointing to a unique communication between neurons and oligodendrocytes prior to the onset of myelination.

With a goal of directly addressing glutamatergic neuronal activity specifically, we used chemogenetics in NexCre mice. Decreased neuronal activity following Gi-DREADD activation resulted in increased OPC differentiation, similar to our UNO model (Figure 2). Electrical and proliferative properties of OPCs at early ages (P6) are indistinguishable across brain regions^43,44^, which is consistent with our data showing comparable OPC responses to altered neuronal activity in both gray matter (cortex) and future white matter (corpus callosum) in these early ages. Further studies using Gq-DREADDs expressed in glutamatergic projection neurons increased neuronal activity, which in turn *inhibited* OPC differentiation into premyelinating and more mature oligodendrocyte states (Figure 3). Again, no difference was noted in cortex relative to corpus callosum OPC responses (Figures 2 and 3). Interestingly, no differences in OPC proliferation occurred in response to increased neuronal activity, as seen in adult models^45–47^, suggesting a developmental context dependent role for neuronal activity in OPC development.

Projection neurons from the cortex cross the midline as early as embryonic day 15, but these neurons are dynamically refined and stabilized during early postnatal development^8,9^. In early development, young neurons are highly exuberant^48^, i.e., over-abundant and highly active, but they are not yet myelinated. Therefore, in early development, oligodendrocytes may respond to these activity cues differently than in later development and adulthood when increased activity promotes myelination. Together, these findings show that across physiological (UNO) and experimentally manipulated contexts (DREADDs), and in multiple brain regions (LOT, somatosensory cortex, and corpus callosum), early developmental neuronal activity normally inhibits oligodendrocyte differentiation and that inhibition of differentiation can be reversed by reducing neuronal activity (Fig. 1-3).

The increases in OPC differentiation were driven by decreased neuronal activity in our models, but we wanted to address the underlying transcriptional changes occurring in early oligodendrocytes in response to reduced neuronal activity. Pathway analysis of gene expression changes in response to decreased neuronal activity unsurprisingly identified increased regulation of gliogenesis, but also identified several interesting pathways related to synapse development and organization. Importantly, we found differential regulation of glutamate receptors: AMPAR subunit gene expression in oligodendrocytes increased in response to reduced neuronal activity, which is comparable to the neuronal response to decreased neuronal activity^49^. This is likely a compensatory response to reduced glutamatergic input to these OPCs upon reduction in neuronal activity, and it highlights the critical role of synaptic signaling in oligodendrocyte development. These analyses also introduce the question of how AMPARs are involved in oligodendrocyte maturation.

OPCs form direct synapses with glutamatergic neurons, and they respond electrically to glutamate through AMPAR stimulation as early as P7 and throughout the lifespan^50^. Numerous studies have investigated the potential role of glutamatergic signaling to oligodendrocytes across development and adulthood^51,52^, including a previous study from our lab showing that the GluA2 subunit of AMPARs regulates OPC migration, and other studies addressing their role in oligodendrocyte survival^21,24^. There is less consensus on the role of AMPARs in oligodendrogenesis. Consistent with the concept that neuronal activity promotes oligodendrogenesis, studies in juvenile and adult mice have shown that AMPAR stimulation also promotes oligodendrogenesis in vivo^23,24^. These studies focused on the role of oligodendroglial AMPARs in juvenile and adult mice, again after the onset of developmental myelination. Other studies have used in vitro and ex vivo models to show that decreased neuronal activity or AMPAR inhibition causes increased OPC proliferation and increased oligodendrocyte differentiation^18–20^. It should be noted that these studies more closely recapitulate an early developmental environment as explants and primary cells are taken from early postnatal mice. Nevertheless, these seemingly conflicting results may indicate a developmental shift in AMPAR-mediated signaling, whereby prior to myelination, glutamatergic activity via AMPARs suppresses oligodendrocyte differentiation, but following the onset of myelination, AMPAR activation facilitates oligodendrogenesis to meet the demand for new myelin. Thus, AMPAR signaling regulates oligodendrocyte development in a stage -and context-dependent manner.

Our ex vivo system allows for greater experimental manipulations to study the effects of neuronal activity and AMPARs. Chemogenetic reduction of neuronal activity increased oligodendrocyte differentiation, recapitulating our in vivo results (Figures 2 and 5), and blocking AMPARs with NBQX also increased oligodendrocyte differentiation, confirming previous studies^18,20^. Excitingly, reduction in neuronal activity with simultaneous AMPAR stimulation reduced the increased differentiation effect of chemogenetically-reduced neuronal activity (Figure 5). These findings suggest that in early development, glutamate released from neurons signals through OPC AMPARs to inhibit oligodendrocyte differentiation. When this inhibition is released by reducing neuronal activity or blocking AMPARs, oligodendrocytes respond by differentiating. Caveats to this interpretation include the fact that Gi-DREADDs only reduce activity from a subset of neurons. Thus bath application of AMPA could be increasing activity of surrounding DREADD-negative neurons, thereby restoring neuronal activity of the tissue, which in this context should reduce differentiation. Additionally, neurons could signal through different glutamate receptors on oligodendrocytes, or through other cells, such as astrocytes, and not act directly on oligodendrocytes.

An exciting result of this experiment is that AMPAR stimulation when neuronal activity was decreased only partially rescued the differentiation phenotype. In this context, the proportion of OPCs was reduced while the proportion of premyelinating cells was greatly increased, and the proportion of mature cells returned to control levels. This suggests that neuronal activity influences oligodendrocytes by multiple mechanisms with AMPAR stimulation regulating the later maturation steps, whereas neuronal activity independent of AMPAR signaling acts to regulate the early phase of OPC differentiation. OPC differentiation to premyelinating cells is still increased with neuronal activity reduction even in the presence of AMPA, but the cells are held at the premyelinating cell stage. These data support the hypothesis that oligodendroglial AMPARs play a key role in communication between excitatory neurons and developing oligodendrocytes, but they highlight the multiple stages at which neuronal activity regulates oligodendrocyte differentiation. Overall, the findings in this study further highlight the context dependence of the interplay of neuronal activity, AMPAR stimulation and oligodendrocyte differentiation. Future studies should focus on the downstream mechanisms of AMPAR signaling in oligodendrocytes.

## Acknowledgements

Wendy Macklin and Won Chan Oh disclose support for the research of this work from (NINDS R21 5R37NS082203-12) and Tessa Allen discloses support from (NINDS F31 1F31NS139593-01A1). This work would not have been possible without the support of the Genomics Core Shared Resource at University of Colorado, Anschutz and their funding from Cancer Center Support Grant (P30CA046934). Finally, we acknowledge Dr. Kennith Jones for his support.

