## Supplemental Figures for "Early developmental neuronal activity inhibits oligodendrocyte differentiation through AMPA receptor activation"

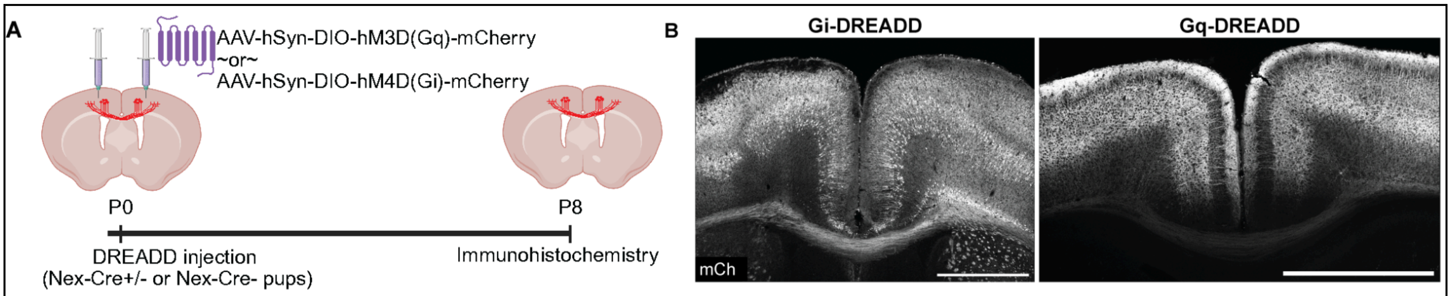


**Supplemental Figure 1: DREADD expression P8.** (A) Experimental design. P0 Nex-Cre+ and Nex-Cre- pups were injected with either Gi-DREADD or Gq-DREADD. (B) Images of the somatosensory cortex and corpus callosum of Nex-Cre+ pups injected with Gi-DREADD (left) and Gq-DREADD (right) showing m-Cherry (mCh) expression. Scale bar is 1mm

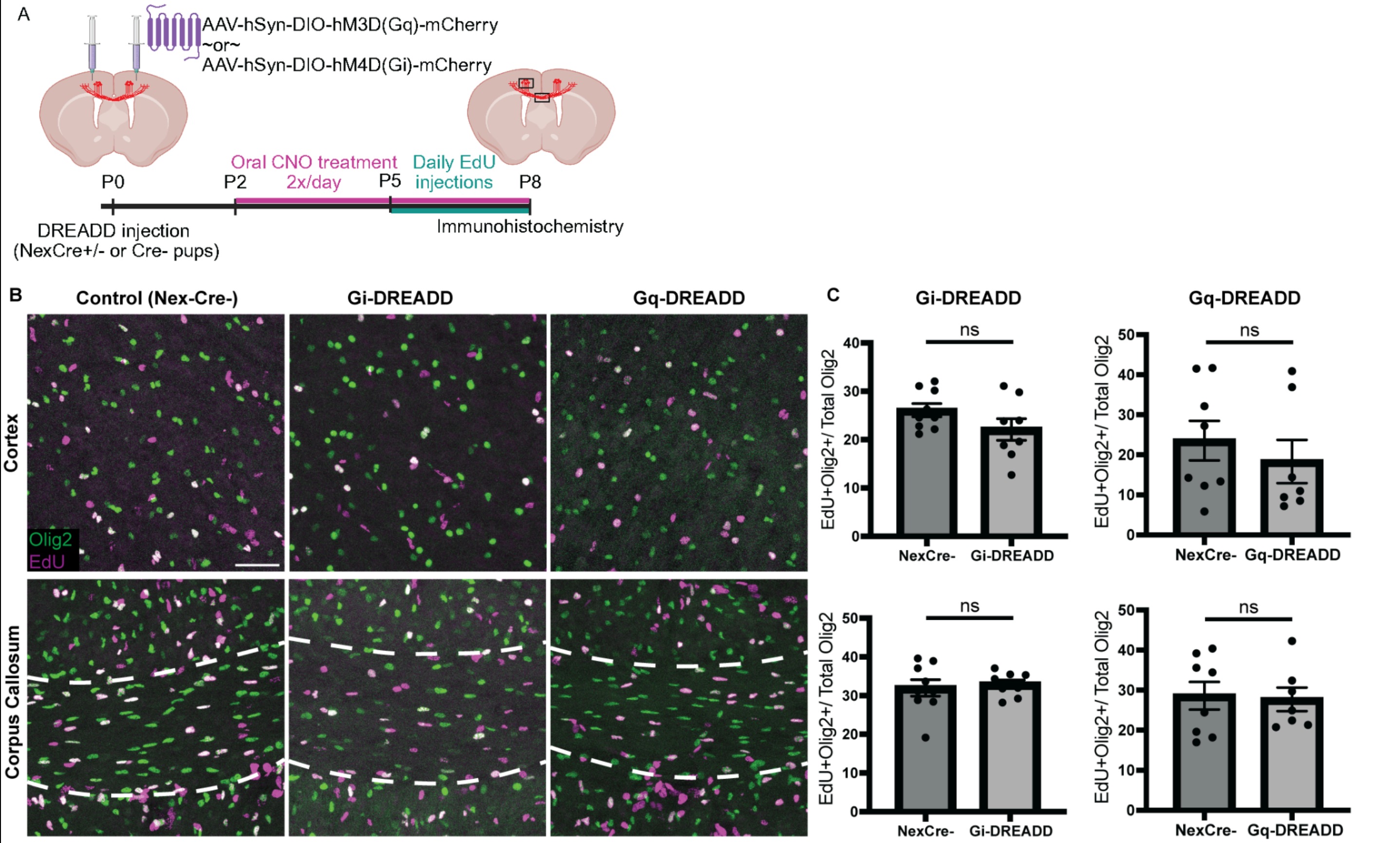


**Supplemental Figure 2: Neuronal activity modulation does not affect OPC proliferation.** (A) Experimental design. P0 or P1 NexCre+ or NexCre- mice were injected bilaterally with Gi-DREADD or Gq-DREADD. CNO was fed to all pups from P2-8. Once daily EdU injections were performed on all pups on P5, P6 and P7. Pups were sacrificed for immunohistochemistry at P8. (B) Images from the cortex (top) and corpus callosum (bottom) showing Olig2 and EdU in NexCre-, Gi-DREADD injected and Gq-DREADD injected from left to right. Scale bar is 50 um (C) quantification of EdU+ Olig2+ cells for Gi-DREADD (left) and Gq-DREADD (right) for the cortex (top) and corpus callosum (bottom). n= 8-9 NexCre- pups, 8 Gi-DREADD pups and 7 Gq-DREADD pups. Student's t-test. ns=not significant

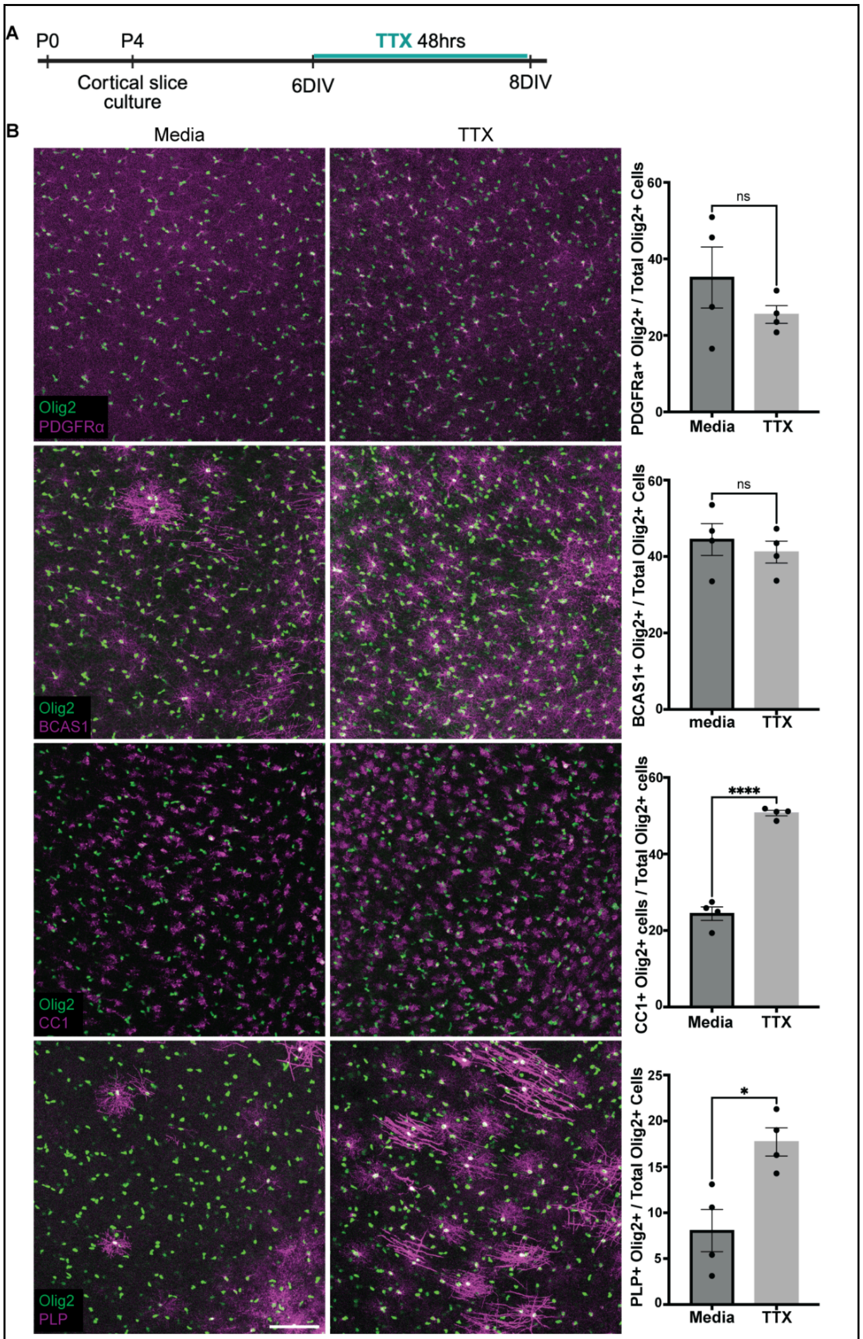


**Supplemental Figure 3: TTX causes increased oligodendrocyte differentiation ex vivo.** (A) Experimental design. P4 WT pups were sacrificed for organotypic cortical slice culture. at 6 days in vitro (DIV), slices were exposed to 1uM TTX for 48hrs. (B) Images and quantification of oligodendrocyte lineage. n= 4 mice. Paired Student's t-test. ns = not significant; * $p<0.01$; **** $p<0.00001$. Scale bar is 50um

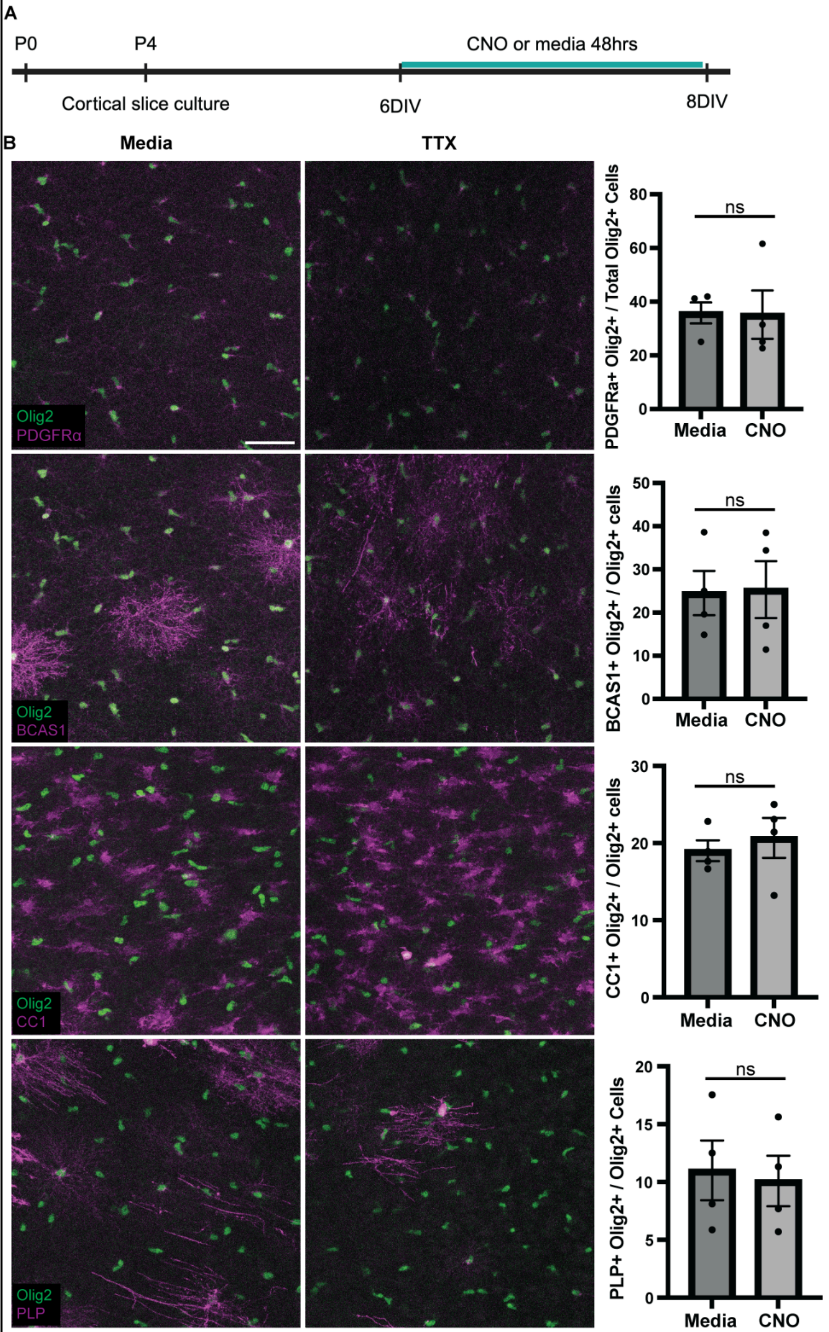


**Supplemental Figure 4: CNO alone does not afftect oligodendrocyte differentiation.** (A) Experimental timeline. Cortical slices were prepared from P4 NexCre+ mice. They were treated for 48hrs from 6-8DIV with CNO or media. (B) Images and quantification of Olig2 along with PDGFRα (OPCs), BCAS1 (premyelinating), CC1 (maturing) and PLP (mature). n=4 animals. Student's t-test. ns = not significant.

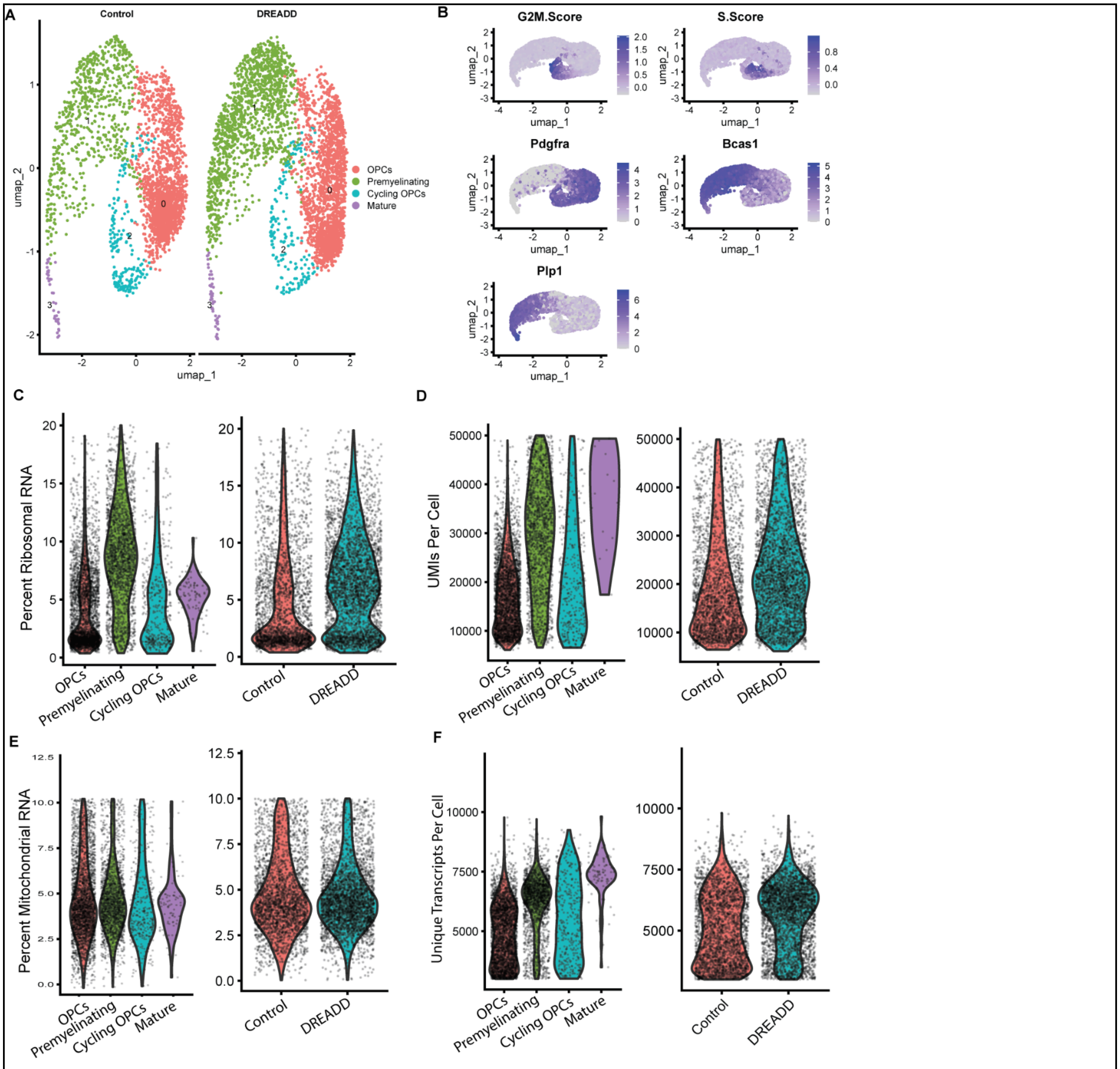


**Supplemental Figure 5. Single cell RNA sequencing quality control data** (A) UMAP embedding showing oligodendrocyte lineage cells from AAV-HM3Di-DREADD injected NexCre+ (n=3 mice pooled) and control NexCre- (n=2 mice pooled) mice all treaed with CNO from p0 to p8. (peach cluster: OPCs, green cluster: premyelinating cells, purple cluster: mature oligodendrocytes, teal cluster: cycling OPCs). (B) Cell cycle and key cluster markers. (C) Percent mitochondrial RNA per cell by cluster (left) and by sample (right). (D) UMIs per cell by cluster (left) and by sample (right). (E) Percent mitochondrial RNA by cluster (left) and by sample (right). (F) Number of unique transcripts detected per cell by custer (left) and condition (right).
